# StayGold Photostability under Different Illumination Modes

**DOI:** 10.1101/2023.11.23.568490

**Authors:** Masahiko Hirano, Yasuo Yonemaru, Satoshi Shimozono, Mayu Sugiyama, Ryoko Ando, Yasushi Okada, Takahiro Fujiwara, Atsushi Miyawaki

**Affiliations:** Laboratory for Cell Function Dynamics, RIKEN Center for Brain Science, 2-1 Hirosawa, Wako-city, Saitama 351-0198, Japan; Biotechnological Optics Research Team, RIKEN Center for Advanced Photonics, 2-1 Hirosawa, Wako-city, Saitama 351-0198, Japan; Evident Corporation, 67-4 Takakura-machi, Hachioji-city, Tokyo 190-0033, Japan; RIKEN CBS-EVIDENT Open Collaboration Center, RIKEN Center for Brain Science, 2-1 Hirosawa, Wako-city, Saitama 351-0198, Japan; Department of Optical Biomedical Science, Institute for Life and Medical Sciences, Kyoto University. Kyoto 606-8507, Japan; Laboratory for Cell Polarity Regulation, RIKEN Center for Biosystems Dynamics Research, Suita, Osaka 565-0874, Japan; Department of Cell Biology, Department of Physics, UBI and WPI-IRCN, The University of Tokyo, Bunkyo-ku, Tokyo 113-0033, Japan; Institute for Integrated Cell-Material Sciences (WPI-iCeMS), Kyoto University, Kyoto 606-8501, Japan; Laboratory of Bioresponse Analysis, Institute for Life and Medical Sciences, Kyoto University. Kyoto 606-8507, Japan

## Abstract

StayGold is a bright fluorescent protein (FP) that is over one order of magnitude more photostable than any of the currently available FPs across the full range of illumination intensities used in widefield microscopy and structured illumination microscopy, the latter of which is a widefield illumination-based technique. To compare the photostability of StayGold under other illumination modes with that of three other green-emitting FPs, namely EGFP, mClover3, and mNeonGreen, we expressed all four FPs as fusions to histone 2B in HeLa cells. Unlike the case of widefield microscopy, the photobleaching behavior of these FPs in laser scanning confocal microscopy (LSCM) is complicated. The outstanding photostability of StayGold observed in multi-beam LSCM was variably attenuated in single-beam LSCM, which produces intermittent and instantaneously strong illumination. We systematically examined the effects of different single-beam LSCM beam-scanning patterns on the photostability of the FPs. This study offers relevant guidelines for researchers who aim to achieve sustainable imaging by resolving problems related to FP photostability.

## Introduction

In many fluorescence imaging experiments, cell samples that contain fluorescent proteins (FPs) targeted to specific organelles are illuminated to excite the FP chromophores. Several illumination modes are available (Fig. 1), and conventional (single-photon excitation) epifluorescence microscopy is categorized into two types. The first type includes widefield (WF) microscopy, which is based on Köhler illumination that involves constant excitation of dyes in a specimen during image acquisition. Conventional WF microscopes are equipped with an arc lamp, whereas recent models use a light-emitting diode (LED) lamp. The typical irradiance (incident flux of radiant energy per unit area) of WF illumination in time-lapse image acquisition with sub-second exposure times is <0.5 W/cm^2^ (refs. 1, 2). On the other hand, structured illumination microscopy (SIM) is a WF illumination-based technique that allows for observation of fluorescent structures at resolutions below the diffraction limit of light^3,4^. SIM experiments that achieve a high spatiotemporal resolution to study the fast dynamics of fine subcellular structures require relatively strong excitation light (1–10 W/cm^2^)^5^. The second type of conventional epifluorescence microscopy includes laser scanning confocal microscopy (LSCM), which employs critical illumination. In single-beam LSCM, the laser beam is focused on single spots sequentially; dyes in the focal plane are excited intermittently but very strongly with an instantaneous irradiance of >1 kW/cm^2^. In multi-beam LSCM, on the other hand, the laser beam is split into approximately one thousand spots and the laser pulse intensity is deconcentrated considerably.

**Figure 1.**
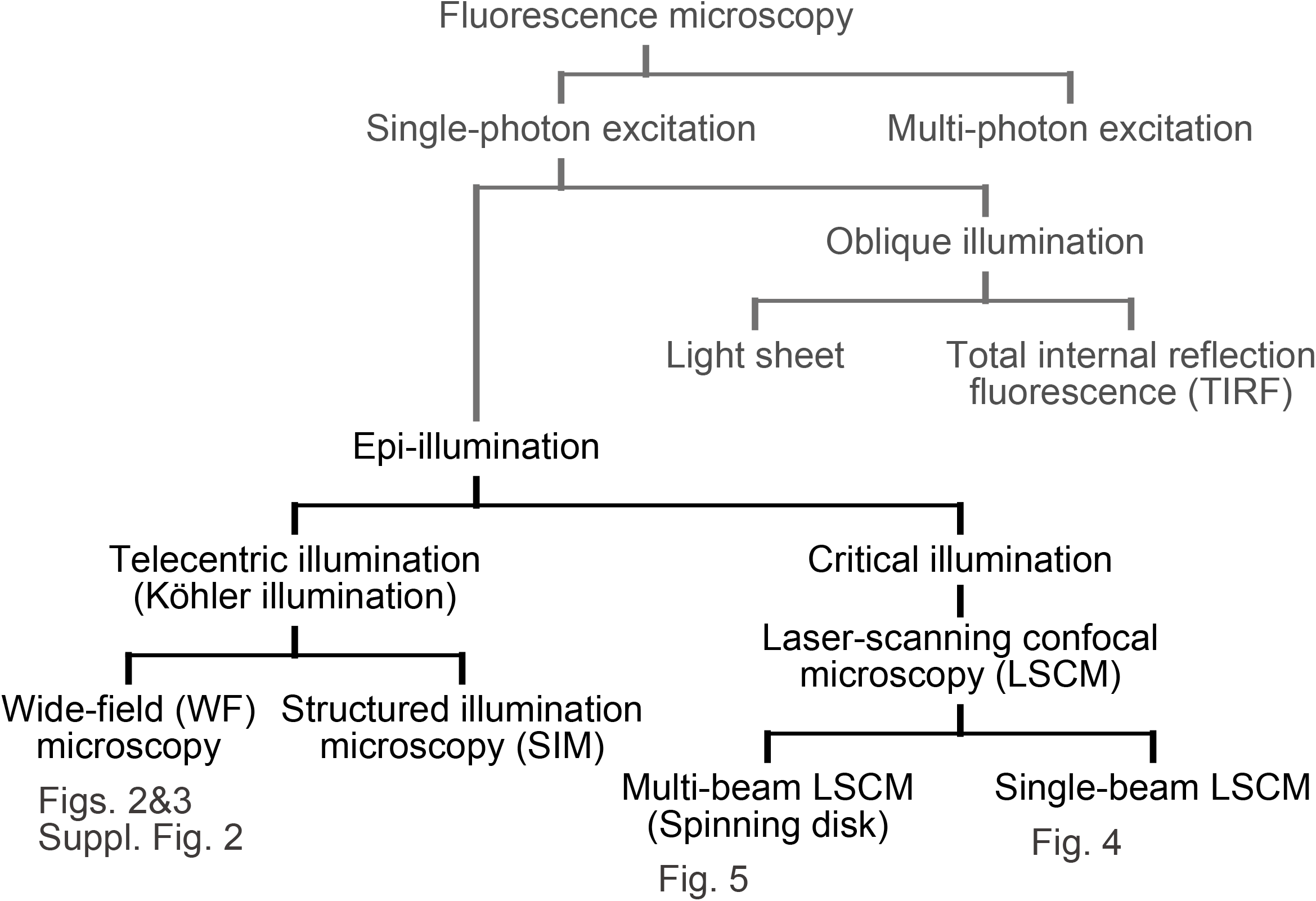
Classification of fluorescence microscopy. Figure # is indicated below the featured microscopy system. The terminology follows ISO 10934:2020, “Microscopes - Vocabulary for light microscopy” (https://www.iso.org/standard/77327.html).

FPs are known for their rapid photobleaching compared with chemical dyes. EGFP is the most classic and popular among useful green-emitting FPs^6^. In the past two decades, a number of FPs have been engineered to be highly efficient in terms of maturation yields. However, these bright FPs, including mClover3 (ref. 7) and mNeonGreen (ref. 8), are less photostable than EGFP. An exception is StayGold, which is derived from the jellyfish *Cytaeis uchidae*^5^. In our previous study, which principally used WF microscopy and SIM (ref. 5), we confirmed that StayGold was not only as bright as mNeonGreen and mClover3 but was also extremely photostable compared with the other FPs (its photostability is more than one order of magnitude higher than that of EGFP). StayGold demonstrates performance that makes sustainable bioimaging possible without being limited by significant photobleaching. For example, this FP enables cell-wide, fast, and continuous super-resolution SIM imaging of endoplasmic reticulum (ER) network dynamics for extended time periods.

In the present study, we compared the practical photostability of StayGold with that of three other green-emitting FPs: EGFP, mClover3, and mNeonGreen. We expressed these four FPs in cultured HeLa cells as fusions to histone 2B (H2B) to be immobilized on the chromatin structures inside the nucleus; this localization was suggested by Shaner et al. for the adequate assessment of FP photostability^9^.

In theory, the decomposition of chromophores (photobleaching) is caused either by interaction with molecular oxygen (O_2_) while the dyes remain in the singlet or triplet excited state, or by strong excitation to higher-order excited states, in which the chromophores are usually more vulnerable to damage. However, it remains unclear how these molecular mechanisms are responsible for the actual FP photobleaching observed in bioimaging experiments. In this regard, it is worthwhile to compare the photostability properties of the extremely photostable FP (StayGold), the most classic FP (EGFP), and the two brightest FPs (mClover3 and mNeonGreen), under different modes of illumination.

## Results

Twenty-four hours after transfection, we comprehensively examined all culture dishes using a low numerical aperture (NA) objective and found that the statistical distribution of brightness was almost the same among cells expressing H2B-StayGold, H2B-EGFP, H2B-mClover3, and H2B-mNeonGreen. From each dish we selected 5–10 cells that were estimated to show the median intensity for the following experiments.

### FP Sensitivity to Chemical Fixation

In the present study, we used fixed as well as live cell samples. To examine beforehand how chemical fixation affects the fluorescence of the four FPs, we performed conventional time-lapse imaging experiments with attenuated arc-lamp illumination (Fig. 2). During the imaging experiments, cells were treated with 4% paraformaldehyde (PFA) at room temperature for 30 min. The fluorescence intensity of StayGold was decreased only slightly, probably because of the change in cell morphology. By contrast, the fluorescence intensities of EGFP and mClover3 quickly dropped down to the background level upon PFA treatment and recovered to near pre-fixation levels after washing with Hank’s Balanced Salt Solution (HBSS). Although the mNeonGreen fluorescence was also sensitive to 4% PFA, both the extinction and recovery processes were very slow, and the fluorescence intensity after recovery was approximately half that of the pre-fixation level (Supplementary Fig. S1).

**Figure 2.**
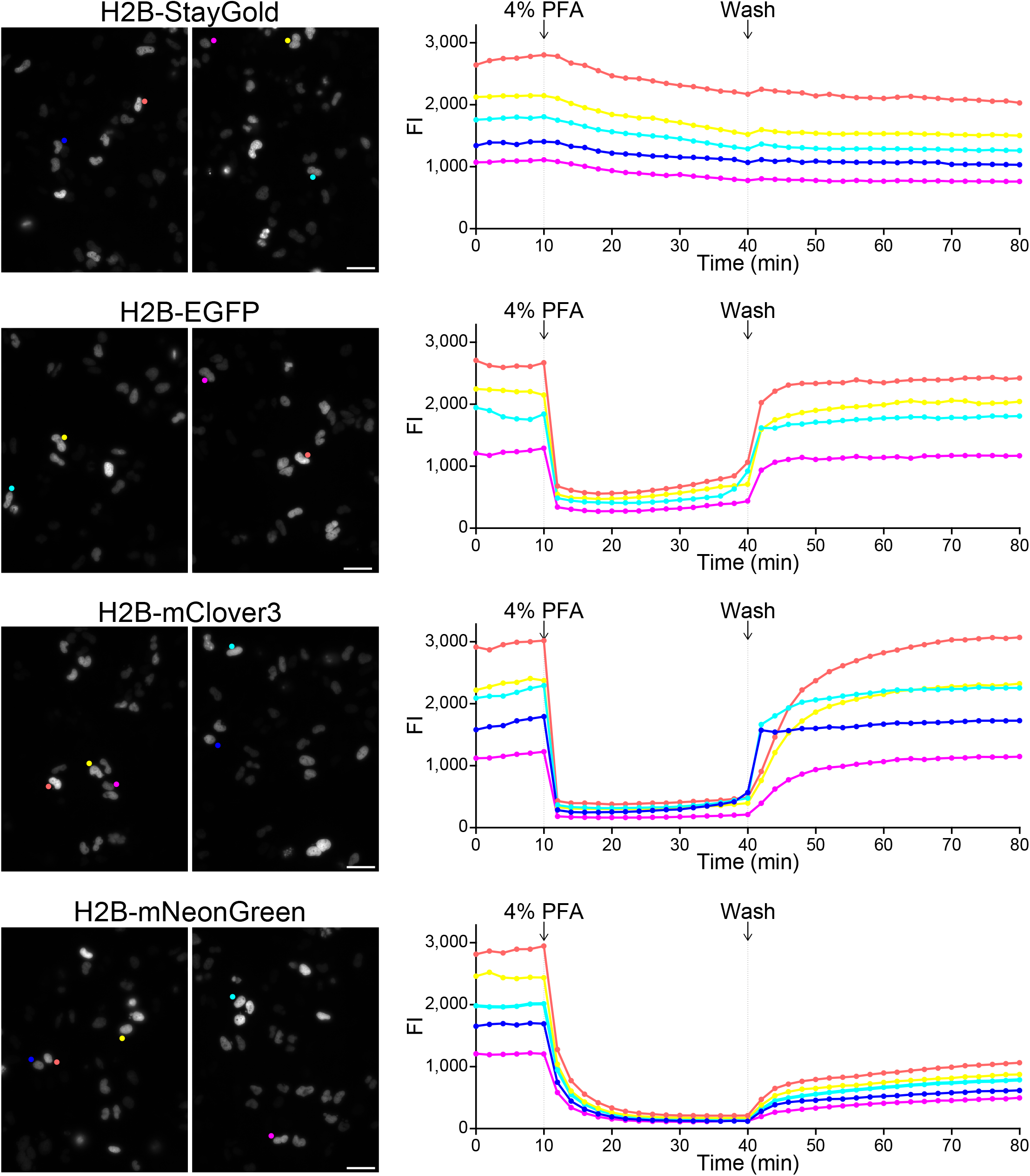
Effects of chemical fixation on the fluorescence of nuclear-targeted FPs. Time-lapse imaging of HeLa cells expressing H2B-StayGold, H2B-EGFP, H2B-mClover3, and H2B-mNeonGreen. *left,* Representative low-magnification fluorescence images before fixation. Scale bars, 50 μm. *right,* Fluorescence intensities (FIs) of several cells are individually plotted against time. Within several hours after washing with HBSS, FIs of recovered fluorescence reached their plateaus. See Supplementary Fig. S1 for the mNeonGreen fluorescence.

### FP Photobleaching under WF Illumination in Live-Cell Samples

We subjected live-cell samples to photobleaching experiments with continuous, unattenuated arc-lamp illumination (Fig. 3A, green lines). We used a 40× objective lens (UPlanSApo 40×/0.95 NA). We measured the radiant flux using a special slide-based power meter (see Methods) and divided it by the view field area to obtain an irradiance of 2.2 W/cm^2^. We adopted the standard method to quantify the photostability of the FPs (ref. 1). Taking into consideration the molecular brightness of each FP (the product of the extinction coefficient at the center wavelength of the illumination (488 nm) and the fluorescence quantum yield of the FP) as well as the irradiance, we plotted the intensity against the normalized total exposure time with an initial emission rate of 1,000 photons/s/molecule (Fig. 3B). The photobleaching half-lives from the initial emission rate of 1,000 photons/s/molecule down to 500 (*t*_1/2_) were 6,847 s for H2B-StayGold, 198 s for H2B-EGFP, 44 s for H2B-mClover3, and 138 s for H2B-mNeonGreen (Table 1).

**Figure 3.**
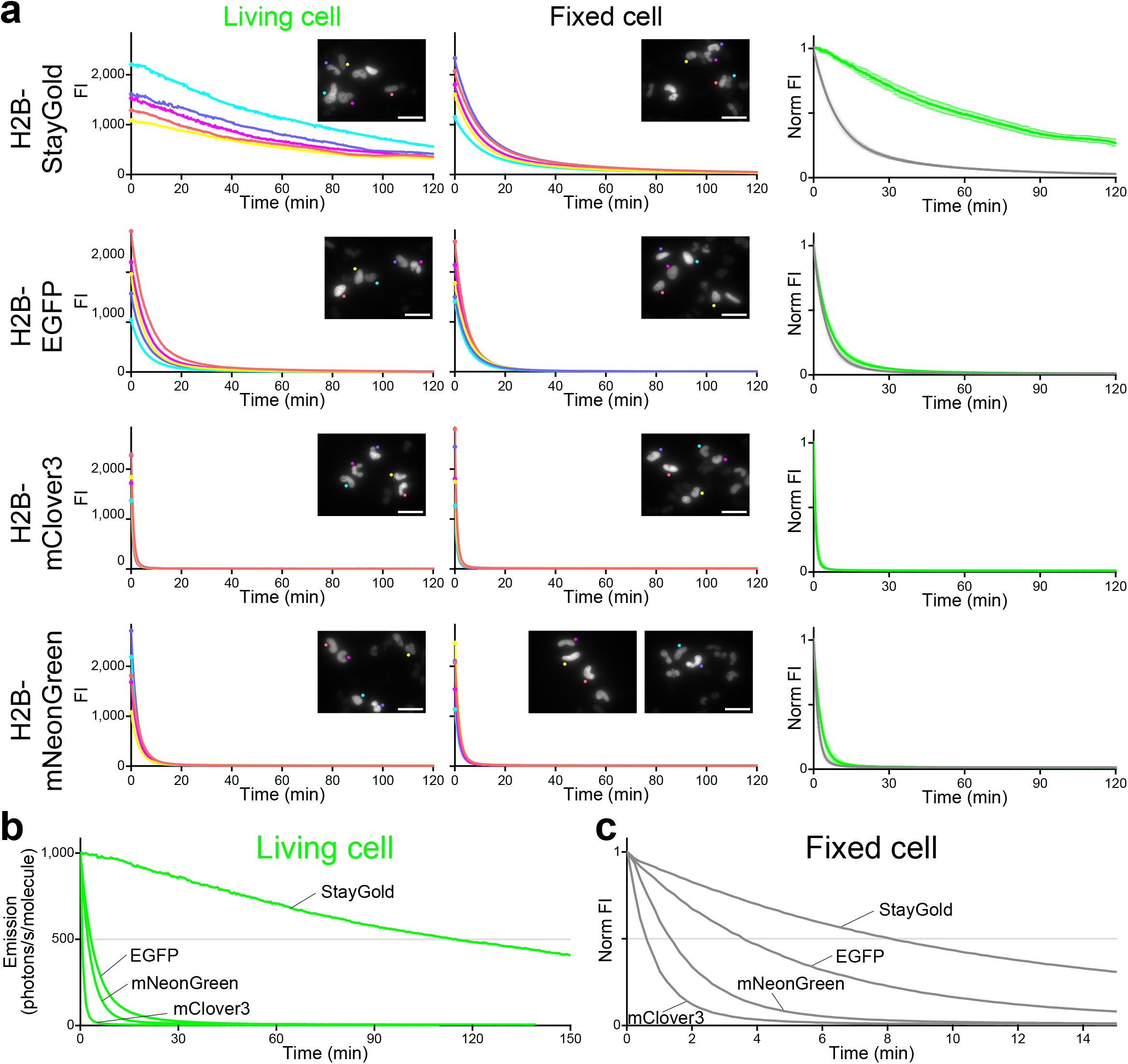
Photostability of StayGold, EGFP, mClover3, and mNeonGreen fused to H2B in HeLa cells under continuous WF (unattenuated arc lamp) illumination. Irradiance: 2.2 W/cm^2^. (A) *left,* Photobleaching curves of the four green-emitting FPs in live-cell samples. *middle,* Photobleaching curves of the four green-emitting FPs in fixed-cell samples. In each photobleaching experiment, five cells were observed (indicated in the first images, *insets*). Scale bars, 50 μm. *right,* Photobleaching curves (green lines: live cells; gray lines: fixed cells) were simply normalized; FI_(t)_/FI_(0)_ was plotted against time. Data points are shown as means ± SD (n = 5 cells). (B) Comparison of the photostability of the four green-emitting FPs in living cells. Photobleaching curves are calculated based on the irradiance and FP molecular brightness (Table 1), plotted as intensity versus normalized total exposure time with an initial emission rate of 1,000 photons/s/molecule. Data points are shown as means (n = 5 cells). (C) Comparison of the photostability of the four green-emitting FPs in fixed cells. As the molecular brightness of FPs in their fixed state has not been determined, simply normalized photobleaching curves shown in (A, *right*) were collected. Data points are shown as means (n = 5 cells).

**Table 1.** Photostability of StayGold and reference green-emitting FPs. The time for photobleaching from an initial emission rate of 1,000 photons/s/molecule down to 500 (*t*_1/2_) in each experiment is shown. All values were measured in this study.

| Illumination |  | Köhler |  | Critical |  |  |  |  |  |
| --- | --- | --- | --- | --- | --- | --- | --- | --- | --- |
| Illumination mode |  | WF |  | Single-beam LSCM |  |  |  | Spinning disk LSCM |  |
| | | | | Transit time (μs) | | | | Rotation speed (rpm) | |
|  |  |  |  | 4.6 | 0.92 | 0.031 | 3.7 | 4,000 | 1,500 |
| Irradiance ($\text{W}/\text{cm}^2$) | | 2.2 | 1.8/1.9 | 0.178 | 0.118 | 0.078 | 0.593 | 1.45 | 1.45 |
| $t_{1/2}$<br>(s) | StayGold | 6,847 | 12,098 | 204 | 145 | 138 | 179 | 1,883 | 1,947 |
|  | EGFP | 198 | 288 | 64 | 70 | 64 | 48 | 327 | 338 |
|  | mClover3 | 44 | 47 | 49 | 45 | 71 | 33 | 72 | 72 |
|  | mNeonGreen | 138 | 167 | 47 | 31 | 46 | 35 | 182 | 209 |
| Related Fig. |  | Fig. 3 | Suppl. Fig. S2 | Fig. 4 |  |  |  | Fig. 5 |  |

We observed the outstanding photostability of StayGold in another experiment that used the H2B-FP fusions (Supplementary Fig. S2, Table 1). Altogether, StayGold was approximately 40, 200, and 50 times more photostable inside the nucleus than EGFP, mClover3, and mNeonGreen, respectively.

### FP Photobleaching under WF Illumination in Fixed-Cell Samples

Next, we examined the photostability in fixed-cell samples under WF illumination. Several hours after fixation and washing with HBSS, the cell samples recovered a substantial fraction of the original fluorescence and were subjected to photobleaching experiments (Fig. 3A, gray lines) under the same optical conditions as those for live cells. Whereas the photobleaching rates of H2B-EGFP, H2B-mClover3, and H2B-mNeonGreen were not significantly altered by fixation, the photostability of H2B-StayGold in the fixed cells was reduced to approximately one-seventh of that seen in live cells. This sensitivity to fixation may suggest that the photostability of StayGold is related to the delicate integrity of the β-barrel. Nevertheless, StayGold was still more photostable after fixation than the other FPs (Fig. 3C).

### FP Photobleaching in Single-Beam LSCM

The photostability of FPs is difficult to evaluate quantitatively in single-beam LSCM^1,9^, whose illumination is intermittent and instantaneously strong. There may be multiple factors that affect the photobleaching rates of FPs in a nonlinear fashion. We discuss some of them below.

First, the illumination point spread function (PSF_ill_) corresponds to the light distribution of the laser that scans the object. The size of the PSF_ill_ is determined solely by the laser wavelength and the NA of the objective lens. On the one hand, the 488-nm laser line is the most appropriate light source for excitation of the four green-emitting FPs. On the other hand, the peak power density is proportional to the square of the objective NA, and we chose the same objective lens as that used in the WF illumination experiments. Accordingly, we employed an Evident FV3000 system equipped with a 40× objective lens (UPlanSApo 40×/0.95 NA) and a 488-nm diode laser; the PSF_ill_ was calculated to have an Airy disk diameter of 0.626 μm (Fig. 4A, solid line).

**Figure 4.**
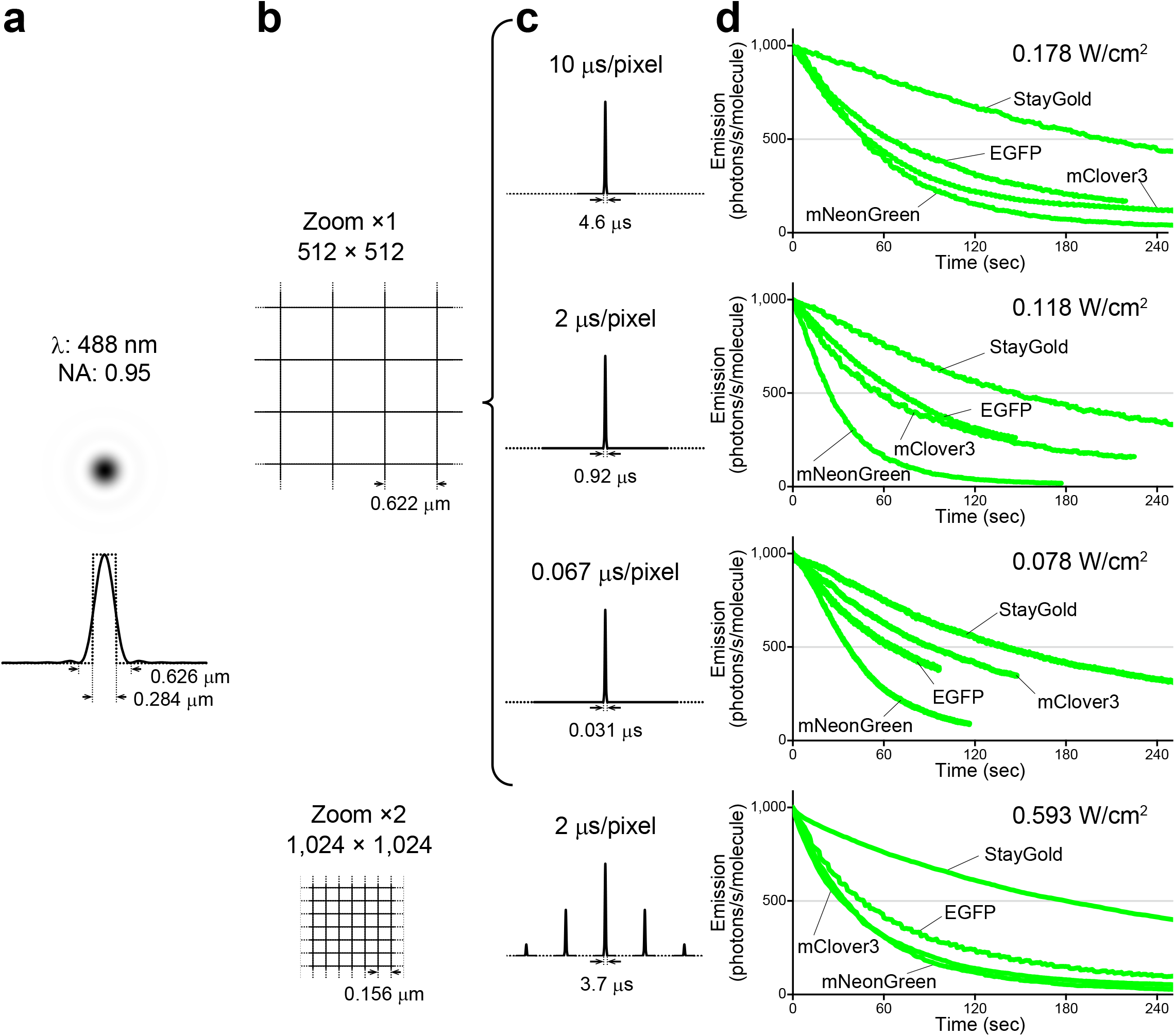
Photostability of StayGold, EGFP, mClover3, and mNeonGreen fused to H2B in HeLa cells with single-beam LSCM. (A) PSF_ill_ in the *xy* plane, calculated with NA = 0.95 and α = 488 nm. The intensity profile (solid line) through the PSF_ill_ center is shown below. The Airy disk diameter is 0.626 μm. The PSF_ill_ section is compared to a square with a side length of 0.284 μm (dotted line). (B) Pixel sizes in the settings of the photobleaching experiments. *top,* zoom factor: ×1; pixel array size: 512 × 512. *bottom,* zoom factor: ×2; pixel array size: 1,024 × 1,024. (C) Scan speeds are indicated above the traces in units of μs/pixel. *top,* Only one illumination pulse was applied during the acquisition of each frame. *bottom,* Multiple illumination pulses were applied during the acquisition of each frame. The pulse widths are indicated below the traces with the calculated transit times. The traces are not drawn to scale. (D) Comparison of the photostability of the four green-emitting FPs in living cells in each experiment using single-beam LSCM. The irradiance value is shown at the upper right. Photobleaching curves are calculated based on the irradiance and FP molecular brightness (Table 1), plotted as intensity versus normalized total exposure time with an initial emission rate of 1,000 photons/s/molecule. Data points are shown as means (n = 5 cells).

Second, there are a variety of beam-scanning patterns. We initially designed a pattern where a given point is not exposed to more than a single illumination pulse during the acquisition of one frame. Using a digital zoom factor of 1× with the aforementioned 40× objective lens, the scanned (imaged) area was 318 μm × 318 μm (= 0.00101 cm^2^). Then, with a pixel array of 512 × 512, the size of each pixel was 0.622 μm (Fig. 4B, top), which was close to the Airy disk diameter (0.626 μm); we understood that this pixel/PSF_ill_ ratio was not ideal for conventional high-resolution imaging. With this configuration, we set the scan speeds at 10 μs/pixel and 2 μs/pixel. We also attempted fast scanning with a speed of 0.067 μs/pixel using an 8-kHz Galvo-Resonant Scanner. Assuming that the PSF_ill_ can be compared to a quadratic prism measuring 0.284 μm on one side (Fig. 4A, dotted line), we calculated the times required for the illumination prism to pass through a given point using these three scan speeds.

The transit times were 4.6 μs, 0.92 μs, and 0.031 μs, respectively. Next, we designed a beam-scanning pattern suitable for conventional high-resolution imaging. Using a zoom factor of 2× and a pixel number of 1,024 × 1,024 with the 40× objective lens, we determined the pixel size of 0.156 μm, which was a quarter of the Airy disk diameter (Fig. 4B, bottom). Then, we set the scan speed at 2 μs/pixel, which was equivalent to a transit time of 3.7 μs.

Under these four different conditions, we continuously illuminated live cells expressing H2B-StayGold, H2B-EGFP, H2B-mClover3, and H2B-mNeonGreen. For each condition, we measured the radiant flux during actual scanning and used it to calculate the scan-averaged irradiance^1^. We generated normalized photobleaching curves (Fig. 4D) and calculated *t*_1/2_ values (Table 1). Remarkably, the *t*_1/2_ of StayGold was dozens of times shorter than that under WF illumination. In addition, we observed a several-fold reduction in *t*_1/2_ for EGFP and mNeonGreen. As a result, although StayGold was still the most photostable of the four green-emitting FPs, the outstanding photostability observed in WF microscopy was attenuated.

Although the pixel dwell time is relative to the pixel size, the transit time should be an absolute value that indicates how long an FP molecule is exposed to light while an illumination pulse passes over it. It seems that a longer transit time accounts for the superior photostability of StayGold compared to the other FPs. In the setting of conventional high-resolution imaging with a transit time of 3.7 μs, for example, StayGold was several times more photostable than any of the other green-emitting FPs (Fig. 4, bottom).

### FP Photobleaching in Multi-Beam LSCM

Our previous study demonstrated that spinning-disk LSCM can optimize the properties of StayGold; we used a Yokogawa CSU X1 equipped with a 100× silicone objective lens (UPlanSApo 100×/1.35 NA) to confirm that a cysteine-less variant of StayGold was more tolerant to continuous illumination than a cysteine-less variant of GFP in the ER lumen^5^. In the present study, we employed an Evident SpinSR10 system equipped with a dedicated objective lens (UPlanXApo 40×/0.95 NA). This objective has the same magnification (40×) and NA (0.95) as the one (UPlanSApo 40×/0.95 NA) used in WF microscopy and single-beam LSCM. Live cells expressing H2B-StayGold, H2B-EGFP, H2B-mClover3, and H2B-mNeonGreen were continuously illuminated with an irradiance of 1.45 W/cm^2^. Although the photostability of the FPs was compared at two disk rotation speeds of 4,000 rpm (maximum) and 1,500 rpm (minimum), we did not find any difference in their photostability. The normalized photobleaching curves and *t*_1/2_ values are shown in Figure 5 and Table 1, respectively. On the whole, StayGold was approximately 6, 27, and 10 times more photostable than EGFP, mClover3, and mNeonGreen, respectively.

**Figure 5.**
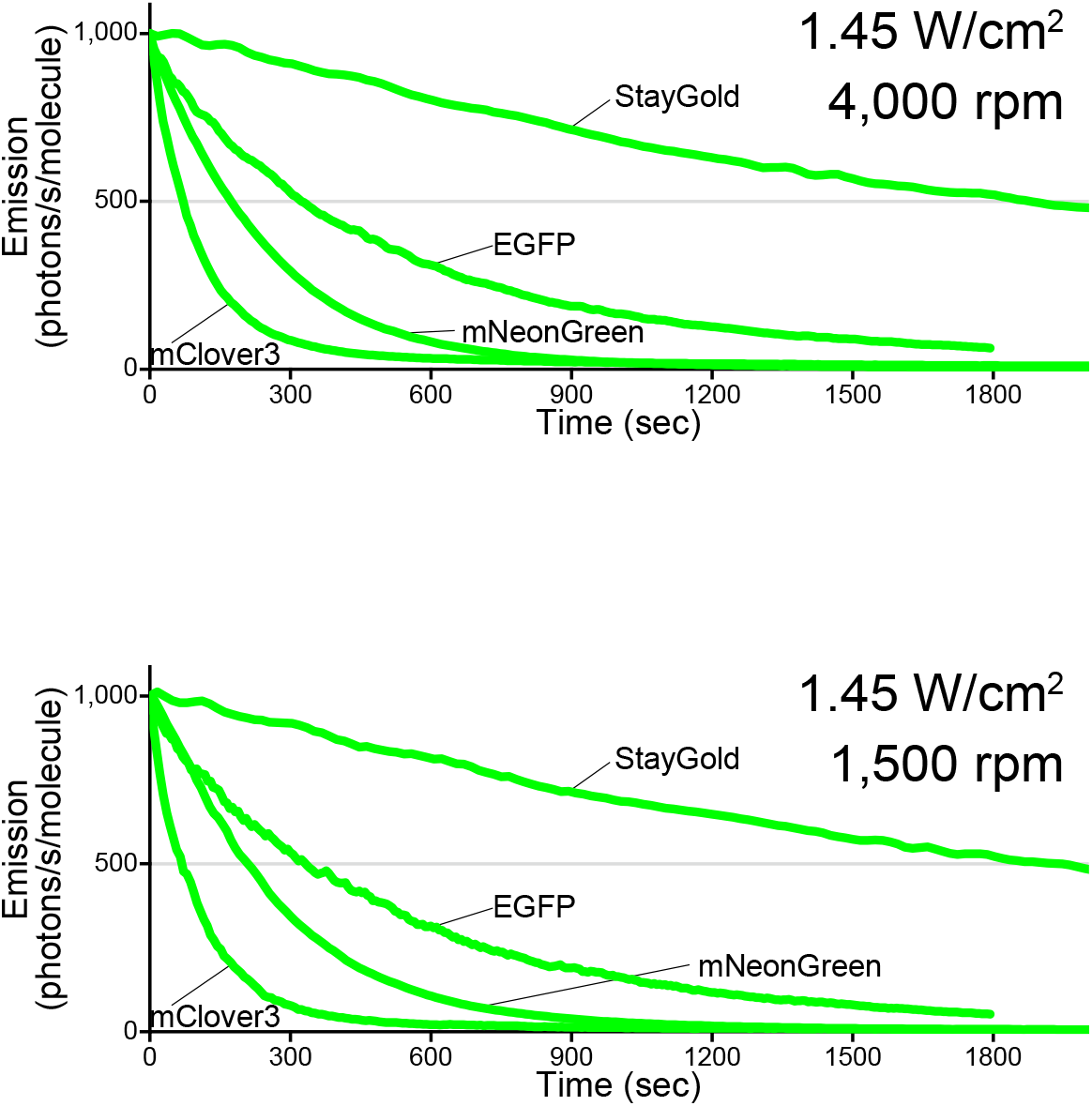
Photostability of StayGold, EGFP, mClover3, and mNeonGreen fused to H2B in HeLa cells with multi-beam LSCM. Comparison of the photostability of the four green-emitting FPs in living cells in each experiment using multi-beam LSCM. The irradiance value and disk rotation speed are shown at the upper right. Photobleaching curves are calculated based on the irradiance and FP molecular brightness (Table 1), plotted as intensity versus normalized total exposure time with an initial emission rate of 1,000 photons/s/molecule. Data points are shown as means (n = 5 cells). The transit time is 3.1–8.7 μs (4,000 rpm) or 8.3–23.2 μs (1,500 rpm).

## Discussion

Among the physical properties of FPs, photobleaching (or photostability) is the most difficult to assess^5^. Shaner et al., who belonged to the laboratory of the late Dr. Roger Tsien, pioneered and successfully standardized the quantification of photostability in the field of FP technology. Assuming that an FP photobleaches with simple exponential decay under continuous illumination, the half-life will be inversely proportional to the rate of the first-order photobleaching reaction. Considering the definitions of the quantum yields for photobleaching and fluorescence, it is apparent that the rate of photobleaching is proportional to the rate of excitation, and eventually to the rate of emission. On the basis of this reciprocity assumption, Shaner et al. established a formulation that the product of the half-life and the rate of emission from a molecule is constant for each FP. The initial emission rate per molecule can be calculated from the irradiance, molar extinction coefficient and fluorescence quantum yield. By substituting this calculated value and the half-life measured from an actual photobleaching curve, it is possible to obtain the time necessary for an FP to photobleach from the initial emission rate of 1,000 photons/s/molecule down to 500. This value (*t*_1/2_) can be used as an absolute index of photostability. Since 2004, Shaner et al. have used this method to quantify the photostability of various FPs principally under the WF mode, i.e., under arc-lamp illumination^10^. Their 2008 report on the development of photostable orange and red FPs (mOrange2 and TagRFP-T, respectively) is the only publication that presents FP photobleaching curves obtained with single-beam LSCM^1^. However, they have suggested that, in principle, the photobleaching efficiency under focused laser illumination is dependent on a number of factors in a nonlinear fashion and is therefore hardly assessable. Compared with single-beam LSCM, by contrast, multi-beam (spinning disk) LSCM produces a dispersed laser illumination and is rather similar to WF microscopy. In this regard, the FP photostability in spinning disk LSCM is fairly quantifiable. In 2017, in fact, Bindels et al. reasonably quantified the photostability of several red-emitting FPs, including mScarlet-I and mScarlet-H variants, using spinning disk LSCM as well as WF microscopy^2^.

In summary, it is difficult to predict the photobleaching behavior of FPs across different microscopy types (illumination modes) and different magnitudes of irradiance. However, with WF illumination this behavior exhibits simple linearity, and WF microscopy has thus been used in most quantitative studies. With the highest NA objective in WF microscopy, unattenuated illumination from an arc lamp or an LED lamp provides an irradiance of up to around 10 W/cm^2^ through a common excitation bandpass filter. In a previous study, we observed the superior photostability of StayGold relative to EGFP across the full range of light intensities (<10 W/cm^2^) of the arc-lamp illumination^5^. The same argument applies to SIM, a WF illumination-based technique. We were able to confirm the outstanding photostability of StayGold in many sustainable SIM imaging experiments using an irradiance of several W/cm^2^ (ref. 5). As demonstrated in the present study, by contrast, the photobleaching behavior of FPs in single-beam LSCM is extremely complicated, and systematic studies are required. For example, whereas mClover3 was always less photostable than mNeonGreen in WF microscopy (Fig. 3, Supplementary Fig. S2) and spinning disk LSCM (Fig. 5), the situation was reversed in single-beam LSCM (Fig. 4D). Moreover, mClover3 photostability was further augmented when using short transit times. We note that the superior photostability of StayGold relative to the other FPs was also decreased by short transit times (Fig. 4D). Such transit-time dependency should be discussed in comparison with the lifetimes of the triplet excited states of these FPs. Another complexity concerning single-beam LSCM is that the scan-averaged irradiance does not necessarily reflect the peak power density that usually exceeds 1,000 W/cm^2^ at an illuminated spot. Such high-intensity illumination will evoke nonlinear effects that cannot be predicted with the current assay system.

Also, the difficulty in the quantitative assessment of photostability stems partly from the lack of a common method for irradiance measurement^5^. The only means for most researchers to assess the photostability of FPs is by performing direct comparisons with reference FPs under the same optical conditions. In the present study, we used a special slide-based power meter that allows for precise measurement of irradiance under all types of illumination modes (see Methods). Low throughput is another reason for the difficulty of quantitative assessment. Continuous illumination naturally precludes parallelization of monitoring, which is why H2B-FP–expressing cells were analyzed sequentially. Furthermore, it always takes a considerable amount of time to monitor the fluorescence of photostable FPs until it fades substantially. Therefore, all the photobleaching experiments in this study were time consuming.

After transfection for FP expression, fixed cells, in addition to living cells, often become observation targets in fluorescence imaging experiments. However, no studies have comparatively investigated the fluorescence properties of FPs before and after chemical fixation. Our current study points out the unique interplay between the photostability of StayGold and its chemical fixation; the fluorescence of StayGold was nearly intact during treatment with 4% PFA, but treated StayGold exhibited a substantial reduction in photostability, which indicates that StayGold is most useful in live-cell imaging. This finding may also shed light on the mechanism underlying the outstanding photostability of this FP. Although the crystal structure of StayGold has been determined to have 1.56 Å resolution^11^, we could not yet fully understand the structural basis of the high photostability. Although future photophysical studies may give us straightforward clues, this report highlights a mystery that StayGold is somehow vulnerable to light from single-beam LSCM, whereas it performs very well in wide-field microscopy, SIM, and spinning-disk LSCM. Thus, the very strong laser illumination used in most photophysical studies to induce photobleaching is not suitable for the adequate analysis of StayGold’s photostability.

## Methods

### Gene construction (nuclear targeting)

The mouse histone 2B (H2B) gene (Fantom3) was amplified using primers containing 5’-*Xho*I and 3’-*Hin*dIII sites, and the restricted product was cloned into the *Xho*I/*Hin*dIII sites of pBS Coupler 1 (ref. 12) to generate pBS Coupler 1/H2B. Also, the green-emitting FP (h-StayGold, EGFP, mClover3, or mNeonGreen) gene was amplified using primers containing 5’-*Bam*HI and 3’-*Xba*I sites, and the restricted product was cloned in frame into the *Bam*HI/*Xba*I sites of pBS Coupler 1/H2B. Lastly, *Xho*I/*Xba*I fragments encoding H2B-(GGGGS)_3_-green-emitting FPs were subcloned into pCSII-EF for transfection. The gene for h-StayGold has mammalian-preferred codons (ref. 5).

### Cell culture, transfection, and fixation

HeLa (HeLa.S3) cells were obtained from American Type Culture Collection (CL-2.2). Cells were cultured on standard 35-mm glass-bottom dishes (3911-035; Iwaki, Japan) in Dulbecco’s modified Eagle’s medium (044-29765; Fuji Film Wako, Osaka, Japan) containing 5% fetal bovine serum (35010107; Corning, Corning, NY) and 1% penicillin/streptomycin (Nacalai Tesque, Kyoto, Japan). The cells were transfected with plasmid DNAs (1.0 μg each) using Lipofectamine 2000 reagent (11668-019; Thermo Fisher Scientific). After washing with HBSS (14025076; Thermo Fisher Scientific), occasionally, the cells were fixed with 4% PFA (P6148; Sigma-Aldrich, St. Louis, MO) in HBSS at room temperature for 30 min.

### WF time-lapse imaging

Cells on 35-mm glass-bottom dishes were incubated in HBSS (14025076; Thermo Fisher Scientific) and imaged on an inverted microscope (IX-81; Evident, Tokyo, Japan) equipped with a standard 75-W xenon lamp, a 20× objective lens (UPlanSApo 20×/0.75 NA), and a cooled charge-coupled device camera (ORCA-AG; Hamamatsu Photonics, Hamamatsu, Japan). A neutral density (ND) filter (12% transmission) (Evident, Tokyo, Japan) was installed in the illuminator. Image acquisition was performed every 2 min with a short exposure time (100–200 ms). The whole system was controlled using AQUACOSMOS software (version 2.6) (Hamamatsu Photonics, Hamamatsu, Japan).

The following optical devices were used. Exciter: 460–480 nm (Evident, Tokyo, Japan) Dichroic mirror: DM485 (Evident, Tokyo, Japan) Emitter: 495–540 nm (Evident, Tokyo, Japan) See Figure 2 and Supplementary Fig. S1.

### WF photobleaching

Living or fixed cells on 35-mm glass-bottom dishes were incubated in HBSS (14025076; Thermo Fisher Scientific) and imaged on an inverted microscope (IX-81; Evident, Tokyo, Japan) equipped with a standard 75-W xenon lamp, a 40× objective lens (UPlanSApo 40×/0.95 NA), and a cooled charge-coupled device camera (ORCA-AG; Hamamatsu Photonics, Hamamatsu, Japan). Whereas no neutral density (ND) filter was installed in the illuminator in principle, an appropriate ND filter was used to attenuate the emitted fluorescence. Image acquisition was performed every 12 s with a short exposure time (300 ms). The whole system was controlled using AQUACOSMOS software (version 2.6) (Hamamatsu Photonics, Hamamatsu, Japan). The area of the illumination field was 0.002375 cm^2^.

The following optical devices were used.

Exciter: 488.0 IF 10 (488 ± 5 nm) (Cheshire Optical)

Dichroic mirror: DM505 (Evident, Tokyo, Japan)

Emitter: BA510IF (> 510 nm) (Evident, Tokyo, Japan) combined with NDX003 (3% transmittance) (Asahi Spectra, Tokyo, Japan)

See Figure 3 and Supplementary Fig. S2.

### Single-beam LSCM photobleaching

Living cells on 35-mm glass-bottom dishes were incubated in HBSS (14025076; Thermo Fisher Scientific) and imaged using an inverted laser scanning confocal microscopy system (FV3000; Evident, Tokyo, Japan) equipped with a 40× objective lens (UPlanSApo 40×/0.95 NA). Green-emitting FPs were excited by a 488-nm diode laser and fluorescence was detected within the wavelength range of 500–600 nm.

See Figure 4.

### Multi-beam LSCM photobleaching

Living cells on 35-mm glass-bottom dishes were incubated in HBSS (14025076; Thermo Fisher Scientific) and imaged on an inverted microscope (IX-83; Evident, Tokyo, Japan) equipped with a 40× objective lens (UPlanXApo 40×/0.95 NA) and a spinning disk field scanning confocal system (pinhole size: 50 μm) (CSU-W1; Yokogawa, Tokyo, Japan). Green-emitting FPs were excited by a 488-nm diode laser and fluorescence was detected within the wavelength range of 500–550 nm. The area of the illumination field was 0.002139 cm^2^.

See Figure 5.

### Measurement of irradiance (W/cm^2^)

The power of excitation light (W) above the objective at the focal plane was measured using a Microscope Slide Power Meter Sensor (S170C; Thorlabs, Newton, NJ) and an Optical Power and Energy Meter (PM100D; Thorlabs, Newton, NJ). The power was divided by the area of the illumination field (cm^2^) to obtain irradiance.

## Acknowledgments

The authors thank K. Higuchi and Y. Ue at RIKEN CBS-Evident Collaboration Center for technical assistance in LSCM; Common Use Equipment in the Support Unit for Bio-Material Analysis, Research Resource Division, RIKEN CBS for technical assistance in multi-beam LSCM; and Dr. M. Kengaku at Kyoto University for continuous support.

This work was supported in part by Grant-in-Aid for Scientific Research (S) (21H05041 to A.M.), Grant-in-Aid for Innovative Areas: Resonance Bio (15H05948 to A.M.) and Information Physics of Living Matters (19H05794, 19H05795 to Y.O.), Japan Science and Technology Agency Core Research for Evolutionary Science and Technology program, “Spatiotemporal dynamics of intracellular components” (JPM JCR20E2 to Y.O.), Marine Biomass Innovation Project (NFRFT-2020-00452 to A.M.), and the Brain Mapping by Integrated Neurotechnologies for Disease Studies from AMED (Brain/MINDS, JP15dm0207001 to A.M.).

## Author Contributions

A.M. conceived the whole study. M.H. performed imaging experiments and analyzed photobleaching of FPs in live and fixed cell samples. A.M., Y.Y. and M.H. designed photobleaching experiments. S.S and R.A. performed gene construction. T.F. and Y.O. analyzed photobleaching of FPs under strong illumination lights. A.M. and M.S. prepared figures. A.M. wrote the manuscript and supervised the project.

