## Supplementary Figs. 1 and 2 for "StayGold Photostability under Different Illumination Modes"

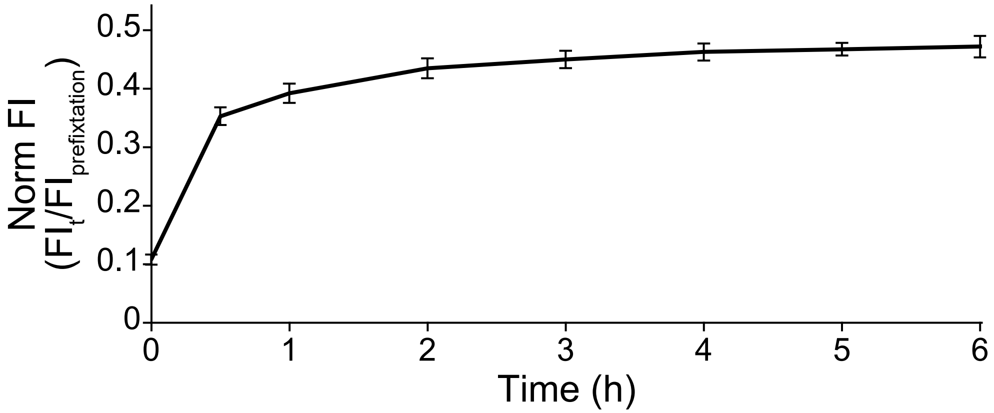


**Supplementary Fig. 1 | Recovery of mNeonGreen fluorescence in fixed HeLa cells after washing with HBSS.**
Data points are shown as means ± SD (n = 5 cells).

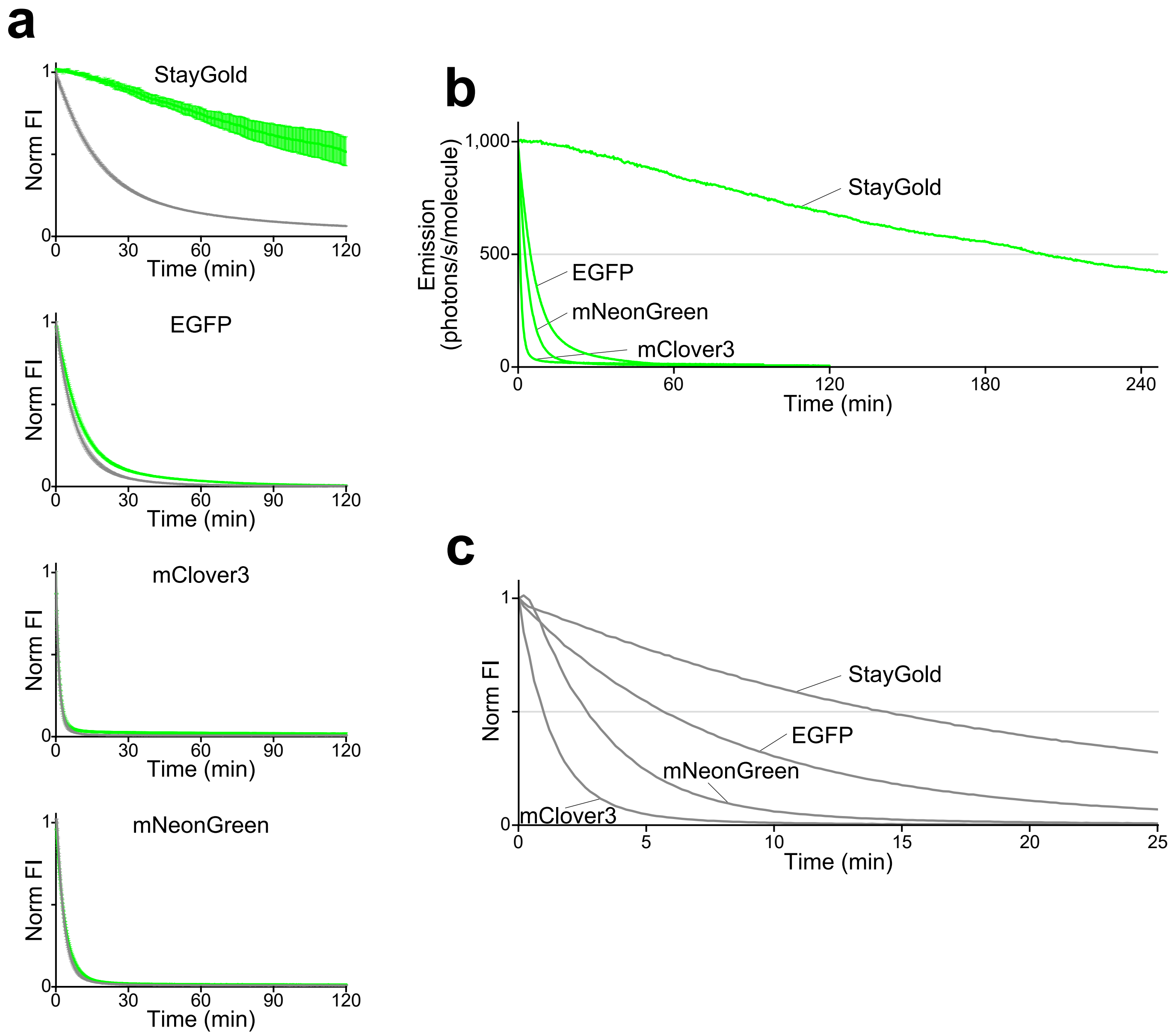


**Supplementary Fig. 2 | Photostability of StayGold, EGFP, mClover3, and mNeonGreen fused to H2B in HeLa cells under continuous WF illumination.**

Irradiance: 1.9 W/cm$^2$ for EGFP, mClover3, and mNeonGreen; 1.8 W/cm$^2$ for StayGold.

(a) Photostability of the four green-emitting FPs in living (green lines) and fixed (gray lines) cell samples. Photobleaching curves were simply normalized; in each experiment, $FI_{(t)}/FI_{(0)}$ was plotted against time.

(b) Photostability in living cells. Photobleaching curves are calculated based on the FP molecular brightness and irradiance (Table 1), plotted as intensity versus normalized total exposure time with an initial emission rate of 1,000 photons/s/molecule. The results were similar to those obtained in our previous WF illumination experiments that monitored the fluorescence of StayGold, EGFP, mClover3, or mNeonGreen distributed throughout the cytosolic and nuclear compartments (ref. 5).

(c) Photostability in fixed cells with an irradiance value of 1.9 W/cm$^2$. The molecular brightness of FPs in their fixed state was not determined. Accordingly, photobleaching curves were based only on irradiance. In each experiment, $FI_{(t)}/FI_{(0)}$ was plotted against time. The coordinate time for StayGold was scaled by 0.947 (=1.8/1.9).
